# A concurrently available negative reinforcer robustly decreases cocaine self-administration in male and female rats

**DOI:** 10.1101/2023.03.29.534800

**Authors:** Madison M. Marcus, Matthew L. Banks

## Abstract

Continued drug-taking despite adverse consequences is hypothesized to be an insidious behavioral hallmark of drug addiction. Although most preclinical research has focused on drug self-administration in the presence of positive punishment, another source of potential adverse consequences is behavioral allocation away from negative reinforcers (i.e., escape/avoid electric shock) and towards drug reinforcers. The goals of the present study were to establish a discrete-trial cocaine-vs-negative reinforcer choice procedure in male and female rats and determine sensitivity of choice behavior to environmental and pharmacological manipulations. Rats could make up to nine discrete choices between an intravenous cocaine infusion (0.32 – 1.8 mg/kg/inf) under a fixed-ratio (FR) 3 schedule and a negative reinforcer (escape or avoidance of electric shock, 0.1 – 0.7 mA) under an FR1 schedule. The negative reinforcer was consistently chosen over all cocaine doses. Lowering shock magnitude decreased negative reinforcer trials, increased omitted trials, and failed to promote behavioral reallocation towards cocaine. Increasing the negative reinforcement response requirement between sessions only increased omitted trials. Introduction of 12-hr extended access cocaine self-administration sessions across two weeks resulted in high daily cocaine intakes but failed to significantly increase cocaine choice. Acute diazepam pretreatment also did not impact choice behavior up to doses that produced behavioral depression. Overall, the lack of behavioral allocation between cocaine infusions and a negative reinforcer suggests these two reinforcers may be economic independents. Additionally, the failure of extended cocaine access to increase cocaine choice highlights the importance of alternative reinforcers and environmental context in preclinical models of drug addiction.

## Introduction

Continued drug use despite adverse consequences is one hallmark characteristic of addiction embedded into multiple diagnostic criteria for substance use disorder. Adverse consequences of cocaine use can take many forms, from social stigma to cardiovascular pain, to overdose death. Cocaine-related overdose deaths have increased every year since 2010, earning cocaine use disorder the title of the “twin” or “silent” epidemic in relation to the current opioid crisis [1–3]. Encounters with the adverse consequences of cocaine use may motivate individuals to seek treatment for cocaine use disorder [4]. Therefore, to improve our fundamental knowledge of the neural mechanisms of cocaine addiction towards the development of candidate treatments, research efforts are increasingly utilizing and developing preclinical procedures that model aspects of this clinical situation.

Addictive drug use despite adverse consequences in nonhumans has commonly been studied by pairing a self-administered intravenous drug infusion with a putative aversive stimulus, such as electric shock. Studies in both rodents and nonhuman primates consistently report that electric shock functions as a positive punisher and decreases cocaine self-administration under a broad range of experimental conditions [5–8]. Of particular interest, some studies report decreased sensitivity of cocaine self-administration to punishment after a history of extended cocaine self-administration, suggesting that extended access cocaine might reveal an “addiction phenotype” of continued drug use despite adverse consequences [9–11]. These previously published studies used single-operant drug self-administration procedures where the primary dependent measure is the rate of behavior and the environmental context includes access to only a single reinforcer (i.e., intravenous drug infusions). The use of single operant drug self-administration procedures poses interpretive complications that can be addressed with preclinical drug choice procedures (for complete review see [12]).

In a typical preclinical drug choice procedure, monkeys or rats have concurrent access to a drug reinforcer and a nondrug positive reinforcer (e.g., food or social interaction) [13–15]. In drug choice procedures, pairing drug-taking behavior with a positive punisher decreases drug self-administration; furthermore, these punishers also promote behavioral reallocation away from the punished reinforcer and towards the unpunished, alternative positive reinforcer [16–19]. In addition to positive reinforcers, negative reinforcers are also available in our natural environment and may also compete with behavior maintained by addictive drugs. In contrast to a positive reinforcer which is operationally defined a stimulus whose *presentation* increases the likelihood of the operant response that preceded it (i.e., cocaine, food), a negative reinforcer is defined as a stimulus whose *removal* increases the likelihood of the operant response that preceded it [20]. Therefore, in addition to electric shock functioning as a positive punisher, studies in both rodents and nonhuman primates have shown that escape or avoidance of electric shock functions as a negative reinforcer [21–23].

The aim of the current study was to develop a novel discrete-trial drug choice procedure in which rats were presented with a conflict: choose negative reinforcement (i.e., foot shock escape or avoidance) and forego a cocaine infusion or forego negative reinforcement and receive both a cocaine infusion and an electric foot shock. There were two main experimental goals. One goal was to conduct a series of parametric studies manipulating independent variables such as reinforcer magnitude and response requirement to determine whether rats would forego escape/avoidance of shock and choose cocaine. In accordance with the cocaine-vs-positive reinforcement literature [24–28], we hypothesized that cocaine-vs-negative reinforcer choice would be sensitive to reinforcer magnitude and response requirement manipulations. The second goal was to determine the effects of extended access cocaine self-administration on cocaine-vs-negative reinforcer choice to test the hypothesis that extended cocaine access altered sensitivity to choose cocaine despite adverse consequences.

## Materials and Methods

### Subjects

A total of 26 Sprague-Dawley rats (13M, 13F; Envigo, Frederick, MD) weighing 230-300g upon arrival were used. Animals were singly housed in a temperature and humidity-controlled vivarium and maintained on a 12-h light/dark cycle (lights off at 6:00 PM). Water and food (Teklad Rat Diet, Envigo) were provided *ad-libitum* in the home cage. Behavioral sessions were conducted five days per week from approximately 11am – 1 pm. Animal maintenance and research were conducted in accordance with the 2011 Guidelines of the National Institutes of Health Committee on Laboratory Animal Resources. Both enrichment and research protocols were approved by the Virginia Commonwealth University Institutional Animal Care and Use Committee.

### Apparatus and Catheter Maintenance

Eight modular operant chambers located in sound-attenuating cubicles (Med Associates, St. Albans, VT) were equipped with electric grid floors (ENV-412 C and ENV-413C) and two retractable levers on the right chamber wall. A set of three LED lights (red, yellow, green) were mounted above the right, drug-associated lever. A white stimulus light was mounted above the left, negative reinforcement-associated lever. Rats were surgically implanted with a custom-made jugular catheter and vascular access port using previously described methods [13] and intravenous (IV) cocaine was delivered by activation of a syringe pump (PHM-100, Med Associates) located inside the sound-attenuating cubicle. Liquid food-maintained responding training sessions occurred in operant chambers equipped with a retractable “dipper” cup (0.1 ml) positioned between the two levers. After each behavioral session, catheters were flushed with gentamicin (0.4 mg), followed by 0.1 ml of heparinized saline (10 U/ml). Catheter patency was verified at the end of each experiment by instantaneous muscle tone loss following IV methohexital (0.5 mg) administration.

### Single Operant Training

Twelve rats (6M, 6F) were initially trained on negative reinforcement (i.e., foot shock escape and avoidance) and 10 rats (5M, 5F) were initially trained on cocaine self-administration. A small group of four (2M, 2F) rats were initially trained on food-maintained responding, then negative reinforcement, and finally cocaine self-administration. Final sample sizes are reported for each experiment.

#### Negative reinforcement training

Rats were initially trained by hand using successive approximation to lever-press to escape electric foot shock during daily 30-min sessions consisting of 60 trials. In each trial, a 3-s foot shock (0.4 mA) was presented along with the left lever and the associated white stimulus light above the lever. Responding was under a fixed-ratio (FR) 1 schedule of reinforcement, such that a single response immediately terminated the shock, retracted the lever, and extinguished the stimulus light. The white house light was illuminated throughout all negative reinforcement training sessions. Acquisition criteria was defined as successful escape of ≥ 80% of the trials for three consecutive days. Once rats met acquisition criteria for escape responding, rats were transitioned to an avoidance-training procedure that also consisted of 60 trials. In this avoidance procedure, shock presentation was preceded by a 30-s avoidance period during which the left lever was extended and associated white stimulus light was on. A single response (FR1) during the avoidance period resulted in cancellation of the upcoming shock for that trial, retraction of the left lever, and extinction of the avoidance stimulus light. If the rat failed to emit a response during the avoidance period, a 3-s shock (0.7 mA) was presented. During shock presentation, the left lever remained extended, and the white stimulus light remained illuminated, signaling the availability of an escape response. The shock intensity was increased to 0.7 mA during avoidance training because pilot studies suggested this shock intensity resulted in the highest rate of acquiring avoidance behavior. Rats were trained on the avoidance procedure for a total of five consecutive days and the number of escape and avoidance trails completed each day were recorded.

#### Cocaine self-administration training

Rats were trained to lever-press for an IV infusion of 0.32 mg/kg cocaine on the right lever under an initial FR1 / 20-s time out schedule of reinforcement during daily 2-hr sessions as previously described [13]. Each session began with a non-contingent cocaine infusion followed by a 60-s time out. The response period was signaled by extension of the right lever and illumination of the associated tricolor stimulus light above the lever. Following each response-requirement completion, the lever retracted, the stimulus light was extinguished, and an IV cocaine infusion was administered. Once rats earned ≥ 30 cocaine infusions during a 2-hr session, the FR requirement was increased to FR3. Acquisition criteria was defined as ≥ 30 cocaine infusions under an FR3 schedule of reinforcement for three days.

#### Food-maintained responding training

Rats were trained to lever press for a 5-s presentation of liquid food (32% Chocolate flavored Ensure™ diluted in water; Abbott Laboratories, Chicago, IL) on the right lever under an initial FR1 / 20-s time-out schedule of reinforcement during daily 2-hr sessions as previously described [13]. Each session began with non-contingent food presentation followed by a 60-s time out. Liquid food availability was signaled by the illumination of the tricolor stimulus light above the right lever. After earning ≥ 30 food reinforcers during a 2-hr session, the FR requirement was increased to FR3. Acquisition criteria was defined as ≥ 30 food reinforcers under an FR3 schedule of reinforcement for three days.

### Cocaine-vs-Negative Reinforcer Choice Procedure

Following successful training of both negative reinforcement- and cocaine-maintained responding alone, rats were trained in the terminal discrete-trial cocaine-vs-negative reinforcer choice procedure. Daily behavioral sessions consisted of two forced trials followed by 9 discrete choice trials. The first forced trial was a cocaine-only trial during which the right, cocaine-associated lever and tricolor stimulus lights were presented. Response requirement (FR3) completion resulted in an IV infusion of the available cocaine dose during the subsequent choice trials. The cocaine forced choice trial incorporated a 30-min limited hold, meaning if the response requirement was not met within 30-min, an IV infusion of the available cocaine dose was administered non-contingently. Following the cocaine-only forced trial, there was a 4 min and 27 s time out during which all stimulus lights were extinguished, and all levers were retracted. Next, a negative reinforcement forced trial was initiated during which the left, negative reinforcer-associated lever was extended, and stimulus light was illuminated for 30-s. Response requirement (FR1) completion on the left lever during this 30-s period resulted in an avoidance response which cancelled the upcoming shock. In the absence of an avoidance response, a 3-s shock stimulus (0.7 mA) was presented and response requirement completion during the shock resulted in an escape response which immediately terminated the shock stimulus. Following completion of both forced trials, choice trials were initiated. During each of the nine choice trials, both the cocaine- and negative reinforcer-associated levers were extended, and the respective stimulus lights were illuminated for 30-s. During this 30-s period, the rat could complete the response requirement on the cocaine-associated lever (FR3) for a cocaine infusion, followed by a 3-s inescapable foot shock (0.7 mA) or the negative reinforcer-associated lever (FR1) to cancel the upcoming shock stimulus (i.e., avoidance response). Response requirement completion on either lever resulted in retraction of both levers, extinction of all stimulus lights, and initiation of a 4 min and 27 s time out period. If the response requirement was not met on either lever during the 30-s avoidance period, a 3-s electric shock (0.7 mA) was presented while both levers remained extended and stimulus lights remained on. During the 3-s electric shock, response requirement completion on the cocaine lever resulted in a cocaine infusion with no shock termination whereas response requirement completion on the negative reinforcer lever resulted in immediate shock termination (i.e., escape response). If the response requirement was not completed on either lever after 3 s, all stimuli including shock were terminated, levers retracted, the trial was recorded as an omission, and a 4 min and 27s time out period was initiated. Rats were tested in the choice procedure five days/week (Mon-Fri).

### Experiment 1: Effect of cocaine dose on cocaine-vs-negative reinforcer choice

In the first experiment, three cocaine doses were tested to determine a cocaine-vs-negative reinforcer choice dose-effect function. Cocaine dose (0.32, 1.0, 1.8 mg/kg/inf) was varied by changing the infusion duration (e.g., 300g rat; 5, 15, 27-s of pump activation, respectively) and cocaine dose presentations were counterbalanced between rats. Each cocaine dose was evaluated over a six-day period. During the first four days, rats were tested at the given cocaine dose vs. negative reinforcer (0.7 mA). On the final two days, the shock stimulus was removed (i.e., no shock condition). The number of days tested under shock and no shock conditions were determined from pilot studies (see Figure S3). Results from the final two days of testing under each condition were averaged and used for data analysis.

### Experiment 2: Effect of shock magnitude on cocaine-vs-negative reinforcer choice

The second experiment systematically determined effects of different shock magnitudes on cocaine-vs-negative reinforcer choice. Based on the results of Experiment 1, 1.8 mg/kg/inf cocaine was used in the choice procedure for experiments 2-5. Shock magnitude was incrementally reduced and then increased every other day (0.7, 0.5, 0.3, 0.1, 0.3, 0.5, 0.7 mA) across 14 test days. Shock intensity was manipulated through custom MedPC programming and verified by daily voltmeter measurements. Results from the second day of testing at a given shock magnitude were reported and used for data analysis.

### Experiment 3: Effect of response requirement on cocaine-vs-negative reinforcer choice

Experiment 3 systematically manipulated the response requirement (1, 2, 4, 8, 16) for the negative reinforcer using a between-day progressive-ratio (PR) schedule for five consecutive days. Shock magnitude (0.7 mA), and cocaine dose (1.8 mg/kg/inf), and cocaine response requirement (FR3) parameters were held constant.

### Experiment 4: Effect of extended cocaine access on cocaine-vs-negative reinforcer choice

Experiment 4 determined the effects of 12-hr extended access cocaine self-administration on cocaine-vs-negative reinforcer choice. Following baseline cocaine-vs-negative reinforcer choice, 12-hr extended access cocaine self-administration sessions were introduced Sunday – Thursday for two consecutive weeks in addition to the daily cocaine-vs-negative reinforcer sessions conducted Monday – Friday. A timeline of this experiment is shown in Figure 4A. Rats were placed in the operant chambers at 6pm and could respond for 0.32 mg/kg/inf cocaine under a FR3 / 10-s time-out schedule of reinforcement with no consequent electric shock. At approximately 6 am, rats were removed from the operant chambers and returned to their home cages. Body weights were assessed daily immediately prior to the choice session. After two weeks of extended cocaine access, a one-week “washout” period occurred wherein only daily cocaine-vs-negative reinforcer choice sessions continued. Data collected on the Friday of this week served as the “post-extended access” data point for subsequent analyses.

### Experiment 5: Effect of acute diazepam treatment on cocaine-vs-negative reinforcer choice

Following a week of baseline cocaine-vs-negative reinforcer choice after Experiment 4, acute diazepam (vehicle, 0.32, 1.0, 3.2, and 10 mg/kg) treatment effects were determined. Presentation order of diazepam doses and vehicle were counterbalanced between rats and administered intraperitoneally ten minutes before the choice session. A one day “washout” session was incorporated between each vehicle or diazepam dose in which no injections were administered but cocaine-vs-negative reinforcer choice sessions still occurred (data not shown).

### Data Analysis

The primary dependent measures in the discrete-trial cocaine-vs-negative reinforcer choice procedure were 1) cocaine trials completed, 2) negative reinforcer (both avoidance and escape) trials completed, and 3) omitted trials. These measures were plotted as a function of cocaine dose or independent variable manipulation. Other dependent measures included the latency to earn a cocaine infusion during the cocaine-only forced trial, the number of cocaine infusions earned during extended-access sessions, and changes in bodyweight relative to pre-extended access baseline. Data were analyzed using repeated-measures one- or two-way analysis of variance, or mixed-effects analysis as appropriate. Sphericity violations were corrected using the Greenhouse-Geisser epsilon. Significant main effects or interactions were followed by planned post-hoc tests that corrected for multiple comparisons. The criterion for significance was set a priori at the 95% level of confidence (p < 0.05), and all analyses were conducted using GraphPad Prism (v 9.4.1, La Jolla, CA).

### Drugs

(-)-Cocaine HCl was provided by the National Institute on Drug Abuse Drug Supply Program (Bethesda, MD, USA). Cocaine was dissolved in sterile water for injection and passed through a 0.22-micron sterile filter before IV administration. Diazepam HCl solution was purchased from a commercial vendor (DASH Pharmaceuticals, Saddle River, NJ). Cocaine and diazepam doses are expressed as the salt form listed above.

## Results

### Cocaine and negative reinforcer training

A total of 21 rats (11 M, 10 F) completed both cocaine and negative reinforcer training and participated in the cocaine-vs-negative reinforcer choice experiments. There was no significant effect of training history on subsequent cocaine-vs-negative reinforcer choice, as shown in Supplemental Table 1 and Figures S1-2. During negative reinforcer training, only a small subset (6/21) of rats acquired avoidance responding (Figure S2).

### Experiment 1: Effects of cocaine dose

Figure 1 shows choice trials completed for cocaine (0.32 – 1.8 mg/kg/inf), negative reinforcement, and trials omitted in the cocaine-vs-negative reinforcer choice procedure under both 0.7 mA shock (i.e., shock) and 0 mA shock (i.e., no shock) conditions. Panels A-C show results from the subset of rats classified as “Avoiders.” Rats were categorized as Avoiders if the subject emitted an avoidance response on at least four trials at each cocaine dose under shock conditions. Rats that emitted an avoidance response on three or fewer trials and instead emitted an escape response were classified as “Escapers” and their results are shown in Panels D-F. Under shock conditions, both avoider and escaper rats completed significantly more trials for the negative reinforcer than cocaine regardless of cocaine dose (Avoiders Trial Type: F(1, 3) = 68.7, p = 0.004; Escapers Trial Type: F(1.3, 21.3) = 47.7, p < 0.001). In the absence of electric shock, there was no significant change in the number of negative reinforcer or cocaine trials completed, nor omissions among Avoiders. In contrast, Escapers were sensitive to removal of electric shock such that cocaine trials increased (Shock Condition: F(1, 16) = 14.9, p = 0.0014) and negative reinforcement trials decreased (Shock Condition: F(1, 16) = 83.8, p < 0.0001). Omitted trials also increased in Escapers during shock removal (Shock Condition: F(1, 16) = 37.5, p < 0.0001).

**Figure 1.**
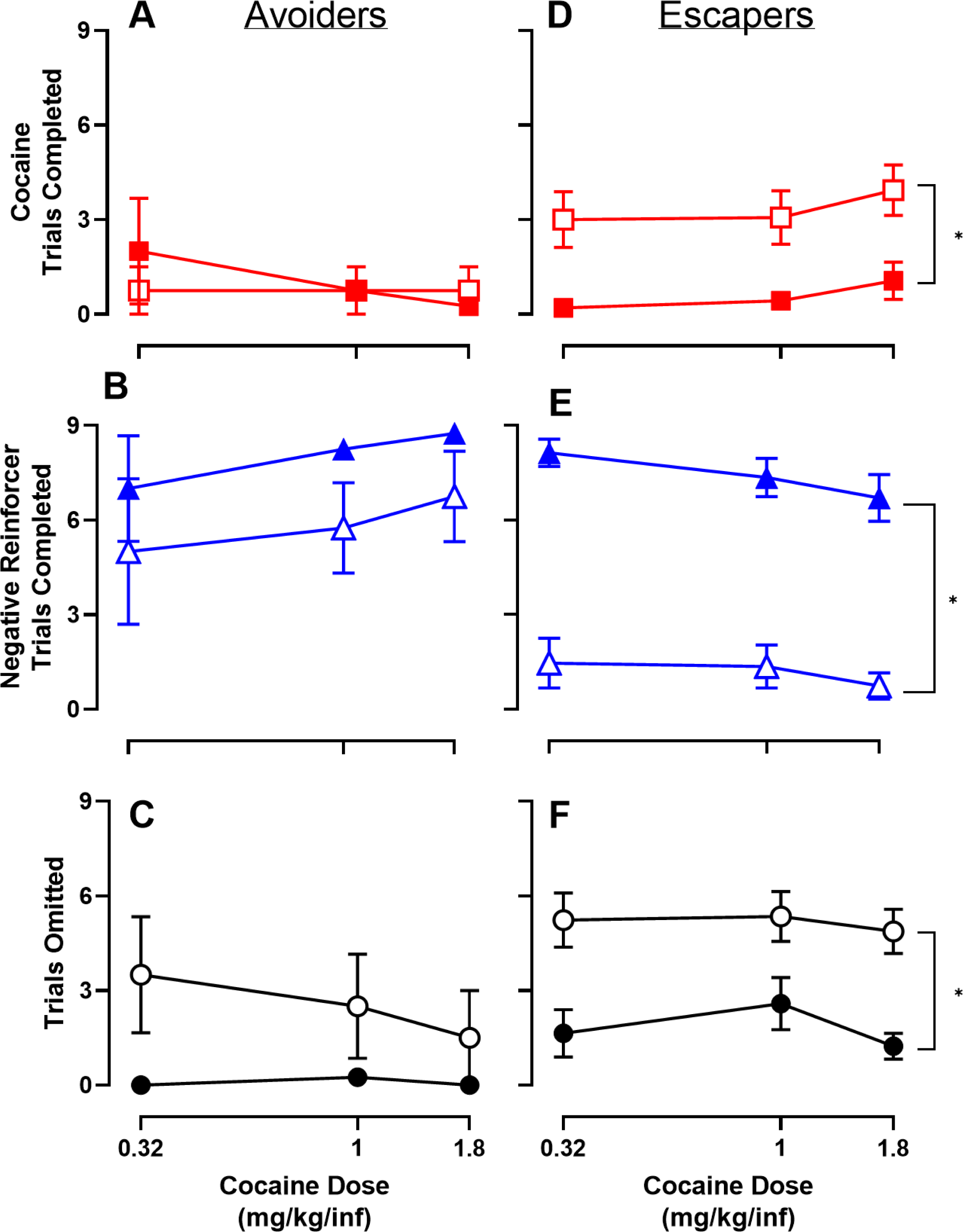
Trials completed for cocaine (FR3), negative reinforcement (foot shock escape or avoidance, FR1), or omitted in a 9 discrete-trial cocaine-vs-negative reinforcement choice procedure as a function of cocaine dose. (A-C) Trials completed by “Avoider” (n = 4, 3F/1M) rats. (D-F) Trials completed by “Escaper” (n = 17, 7F/10M) rats. Filled symbols denote shock (0.7 mA) condition; open symbols denote no shock condition. All points represent mean ± SEM. Brackets represent significant main effect of shock condition. “Avoider/Escaper” classification: under shock conditions, “avoider” rats avoided at least 4 of the 9 trials at each cocaine dose tested. “Escaper” rats avoided 3 or fewer trials at each cocaine dose.

### Experiment 2: Effects of shock magnitude

Figure 2 shows the effect of manipulating shock magnitude on behavioral allocation between 1.8 mg/kg/inf cocaine and negative reinforcement. Rats were classified as Avoiders if at least four trials were avoided at each shock amplitude tested and classified as Escapers if three or fewer trials were avoided, and escape responses were emitted. Results from Avoiders revealed a significant main effect of trial type (Trial Type: F(1.1, 5.7) = 9947, p < 0.0001) and a post-hoc analysis correcting for multiple comparisons revealed that Avoiders completed significantly more trials for the negative reinforcer compared to cocaine across all shock intensities; (F(1.1, 5.7) = 12864, p < 0.001; Fig 2A). The number of omitted trials was low across all shock intensities in Avoiders. In contrast, negative reinforcement trials completed in Escapers was sensitive to shock magnitude (Trial Type: F(1.6, 17.1) = 9.8, p = 0.003; Trial Type × Shock Magnitude: F(4.1, 40.7) = 12.3, p < 0.0001; Fig 2B). At shock magnitudes ≥ 0.5 mA, significantly more negative reinforcer trials were completed over omitted and cocaine trials. At the smallest shock intensity (0.1 mA), more trials were omitted than completed for the negative reinforcer. There were never significantly more cocaine trials completed than negative reinforcer trials completed at any shock amplitude.

**Figure 2.**
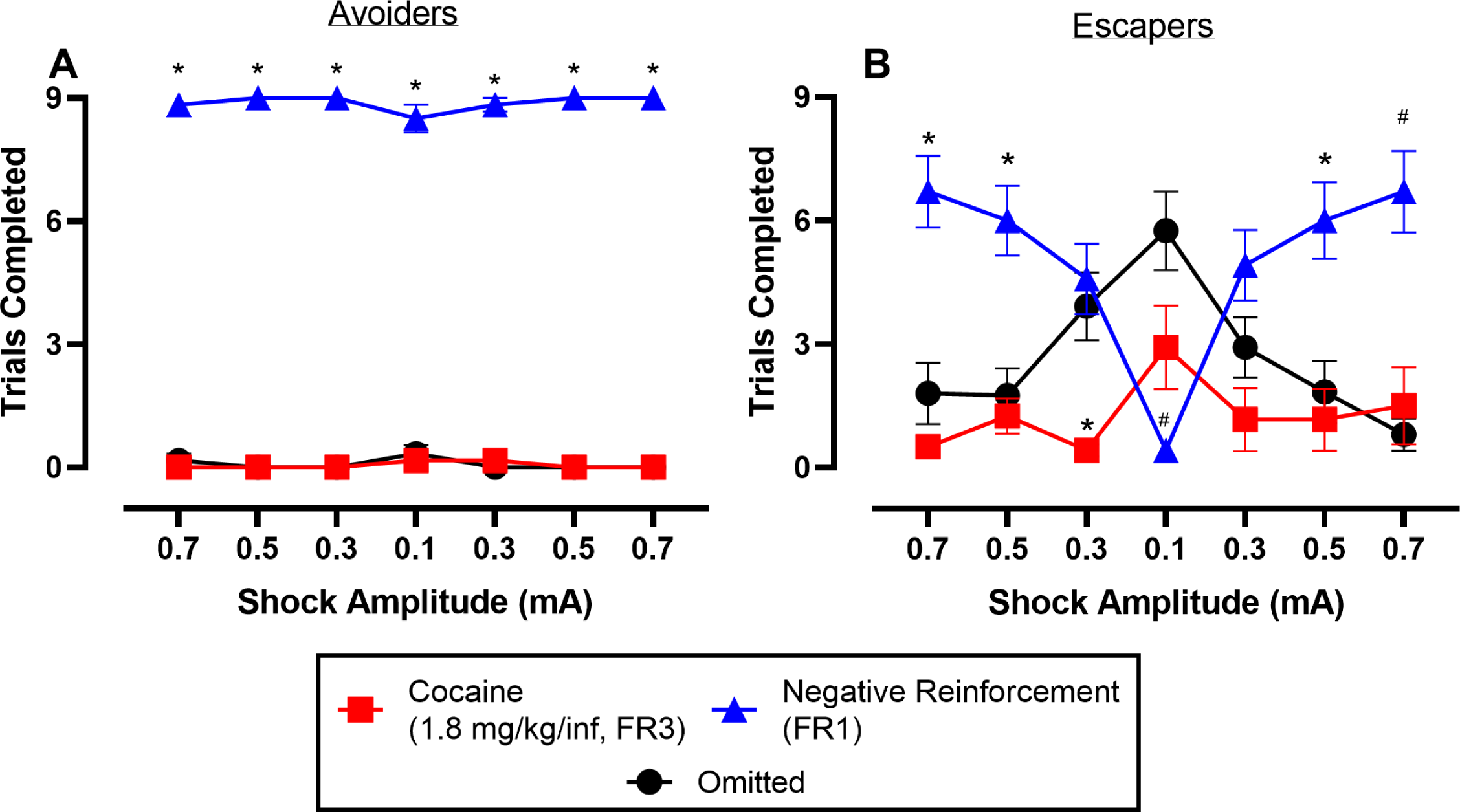
Effect of shock magnitude on cocaine-vs-negative reinforcer choice. Abscissae: shock magnitude. Ordinates: number of trials completed for cocaine, negative reinforcement, or omitted in the 9 discrete-trial choice procedure. All points represent mean ± SEM from the second test day. (A) trials completed by “Avoider” (n = 6, 4F/2M) rats; *significance (p < 0.05) compared to both cocaine/omitted trials. (B) trials completed by “Escaper” (n = 12, 4F/8M) rats; symbols denote significant (p < 0.05) comparisons within a shock amplitude: *different from other two reinforcers, ^#^different from omissions only; “Avoider/Escaper” classification: under shock conditions, “avoider” rats avoided at least 4 of the 9 trials at each shock intensity tested. “Escaper” rats avoided 3 or fewer trials at each shock intensity.

### Experiment 3: Effects of response requirement

Figure 3 shows effects of increasing the response requirement for the negative reinforcer on cocaine-vs-negative reinforcer choice. Increasing response requirements for the negative reinforcer resulted in decreased negative reinforcer trials completed and increased omitted trials (Response Requirement × Trial Type: F(3.3, 28.1) = 16, p < 0.0001). Cocaine trials were unaltered by manipulating the negative reinforcer response requirement. As an additional experiment, choice behavior was determined when the response requirement for both the negative reinforcer and 1.8 mg/kg/inf cocaine were equal at FR1 for five consecutive days. Figure S4 shows that trials completed for cocaine, negative reinforcement, or omitted were not different when the cocaine response requirement was either FR3 or FR1.

**Figure 3.**
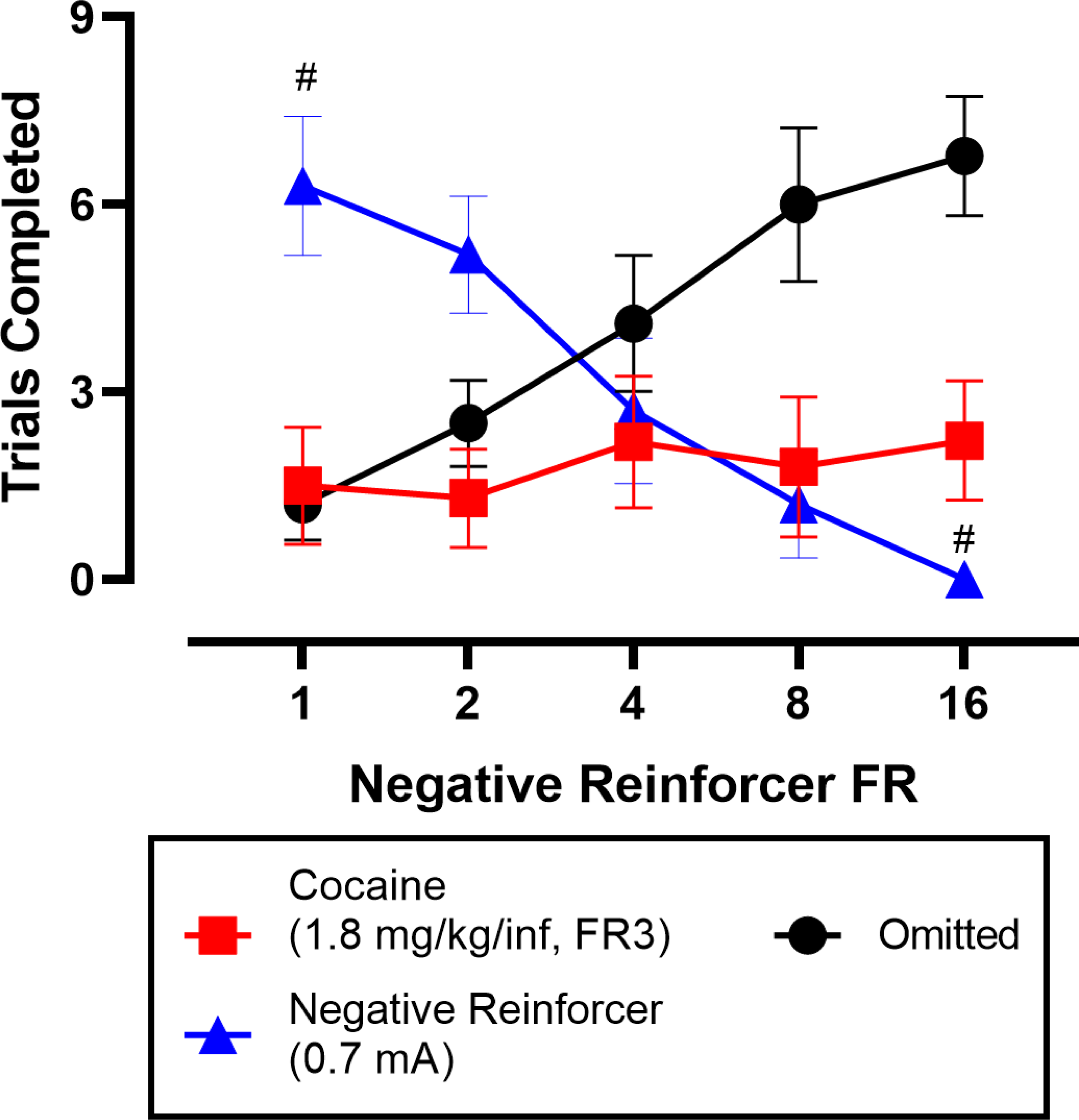
Effect of negative reinforcer response requirement on cocaine-vs-negative reinforcer choice. All points represent the mean ± SEM. ^#^denotes significant (p < 0.05) differences from omitted trials; n = 10 (6F / 4M).

### Experiment 4: Effects of extended cocaine access

Effects of 12-hr extended access cocaine (0.32 mg/kg/inf, FR3 / 10-s time out) self-administration were determined on cocaine-vs-negative reinforcer choice in two cohorts that varied in shock magnitude (0.7 vs. 0.3 mA; Fig. 4A). Cocaine daily intake during the extended access sessions was approximately 96 mg/kg/day (Fig 4B) and only a significant decrease in cocaine infusions was observed on Day 3 compared to Day 1 in the 0.7mA shock intensity group (F(3.4, 32.5) = 9.3, p < 0.0001; Fig 4B). Bodyweight also decreased from baseline in both shock intensity groups (0.7mA Shock Condition: F(2.9, 28.3) = 13.5, p <0.0001; 0.3 mA Shock Condition: F(1.9, 13.5) = 5.9, p = 0.01; Fig 4C). Extended access cocaine did not significantly alter cocaine trials completed under either the 0.7 or 0.3 mA shock condition (Fig 4D). More trials were completed for the negative reinforcer under the 0.7 mA shock condition compared to the 0.3 mA shock condition and there was no significant effect of extended cocaine access (Shock Condition: F(1, 11) = 12, p = 0.005; Fig 4E). There were more omitted trials in the 0.3 mA shock condition compared to the 0.7 mA shock condition (Shock Condition: F(1, 11) = 15.6, p = 0.002; Fig 4F). Although there was a main effect of experimental day on omitted trials (Experimental Day: F(4, 43.9) = 2.9, p = 0.033), post-hoc analysis correcting for multiple comparisons did not report significant changes from baseline omissions in either shock intensity group. In addition, no sex differences were observed (Figures S5-6). Individual subject analysis reveals that of the 20 rats tested, only one increased cocaine choice following extended cocaine access (Figure S7).

**Figure 4.**
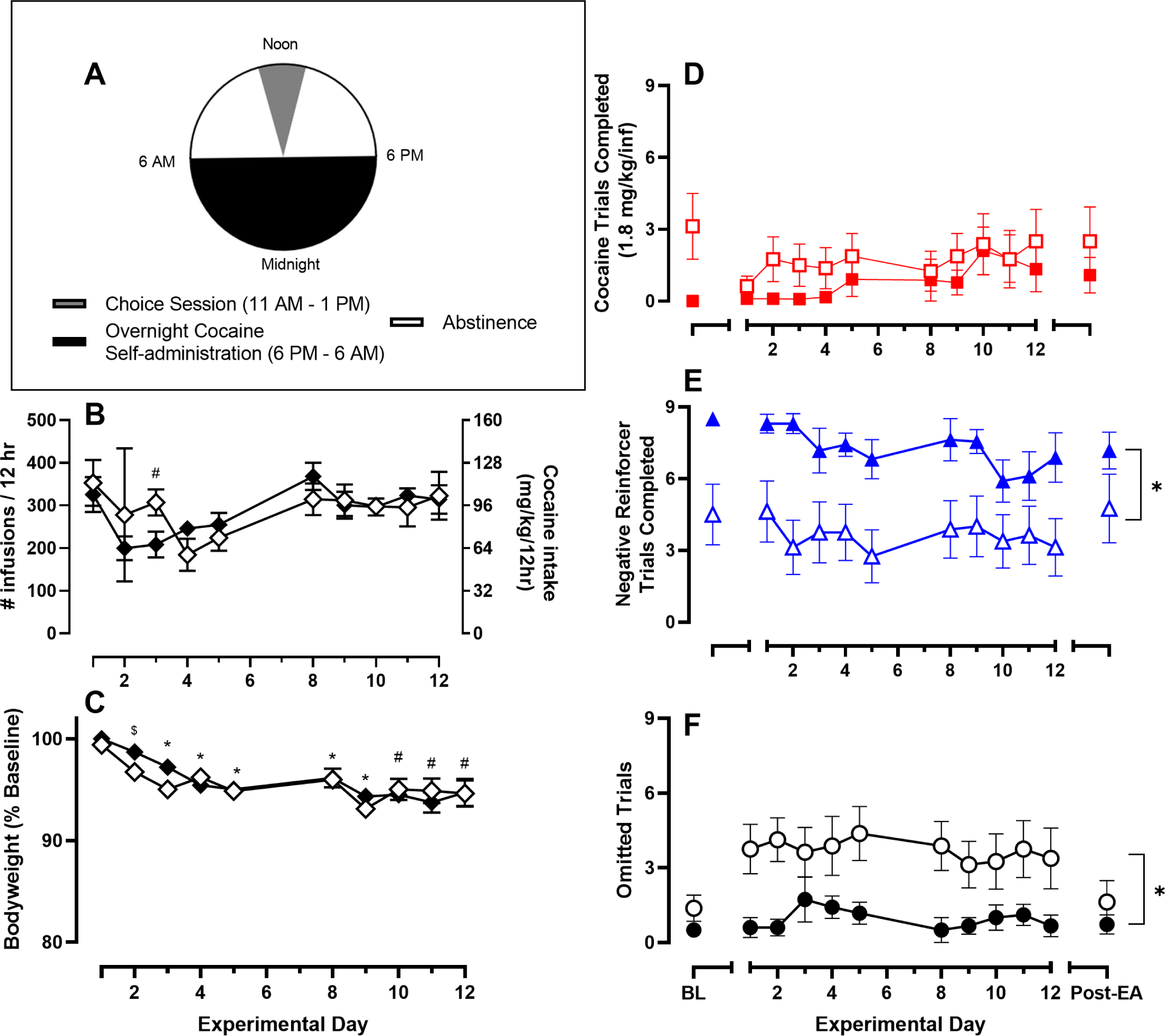
Effect of extended cocaine access on overnight cocaine intake, bodyweight, and choice trials completed for cocaine, negative reinforcement, or omitted. Abscissae: experimental day. (A) schematic of experimental design (B) number of cocaine infusions earned (FR3 / TO10, 0.32 mg/kg/inf) during the overnight session (C) change in bodyweight expressed as a percentage of baseline. (D – F) Trials completed for cocaine, negative reinforcer, or omitted during the choice session. Filled and open circles denote rats tested in the cocaine-vs-negative reinforcer choice procedure under 0.7mA (n = 12, 6F/6M), or 0.3mA (n = 8, 4F/4M) shock condition, respectively. Baseline (BL) is the Friday prior to initiating extended access (EA) cocaine on the following Sunday. Post-EA is 7 days after terminating extended access sessions. All points represent the mean ± SEM. Brackets represent significant main effect of shock condition. ^#^ denotes significant (p < 0.05) differences from Day 1 only in the 0.7mA shock group; ^$^different from Day 1 only in the 0.3mA shock group. *different from Day 1 in both shock intensity groups.

### Experiment 5: Effects of acute diazepam treatment

Figure 5 shows acute diazepam effects on cocaine-vs-negative reinforcer choice under both 0.7 and 0.3 mA shock conditions. More cocaine trials were completed under 0.3 than 0.7 mA shock conditions (Shock Condition: F(1, 6) = 8.1, p = 0.029; Fig 5A). Although there was a main effect of diazepam dose (Dose: F(1.8, 10.7) = 6.1, p = 0.02), diazepam did not alter cocaine trials completed upon post-hoc analysis that corrected for multiple comparisons. More trials were completed for the negative reinforcer under the 0.7 mA shock condition than the 0.3 mA shock condition (Shock Condition: F(1, 6) = 13.3, p = 0.011; Fig 5B). There was a main effect of diazepam dose (Dose: F(1.4, 8.3) = 18.9, p = 0.001) and 10 mg/kg diazepam decreased the number of negative reinforcer trials completed under 0.7mA shock condition (F(2.2, 12.7) = 26.4, p <0.0001; Fig 5B). More trials were omitted in the 0.3 than 0.7 mA shock condition (Shock Condition: F(1, 6) = 8.9, p = 0.025; Fig 5C). There was a main effect of diazepam dose (Dose: F(1.9, 11.5) = 108.6, p < 0.0001), and 10 mg/kg diazepam increased omissions under both shock conditions (0.7mA Shock Condition: F(1.4, 7.8) = 115.2, p <0.0001; 0.3mA Shock Condition: F(2.2, 9.0) = 29.75, p < 0.0001). Furthermore, there was a main effect of diazepam dose on latency to complete the FR3 response requirement for a cocaine infusion during the initial forced trial component (Dose: F(2.6, 15.9) = 25.9, p < 0.001), such that the start latency was significantly increased relative to vehicle after 3.2 and 10 mg/kg diazepam in the 0.7 mA shock cohort (F(1.7, 11.7) = 69.9, p < 0.001; Fig 5D). All rats reached the 30-min limited hold at the highest diazepam dose.

**Figure 5.**
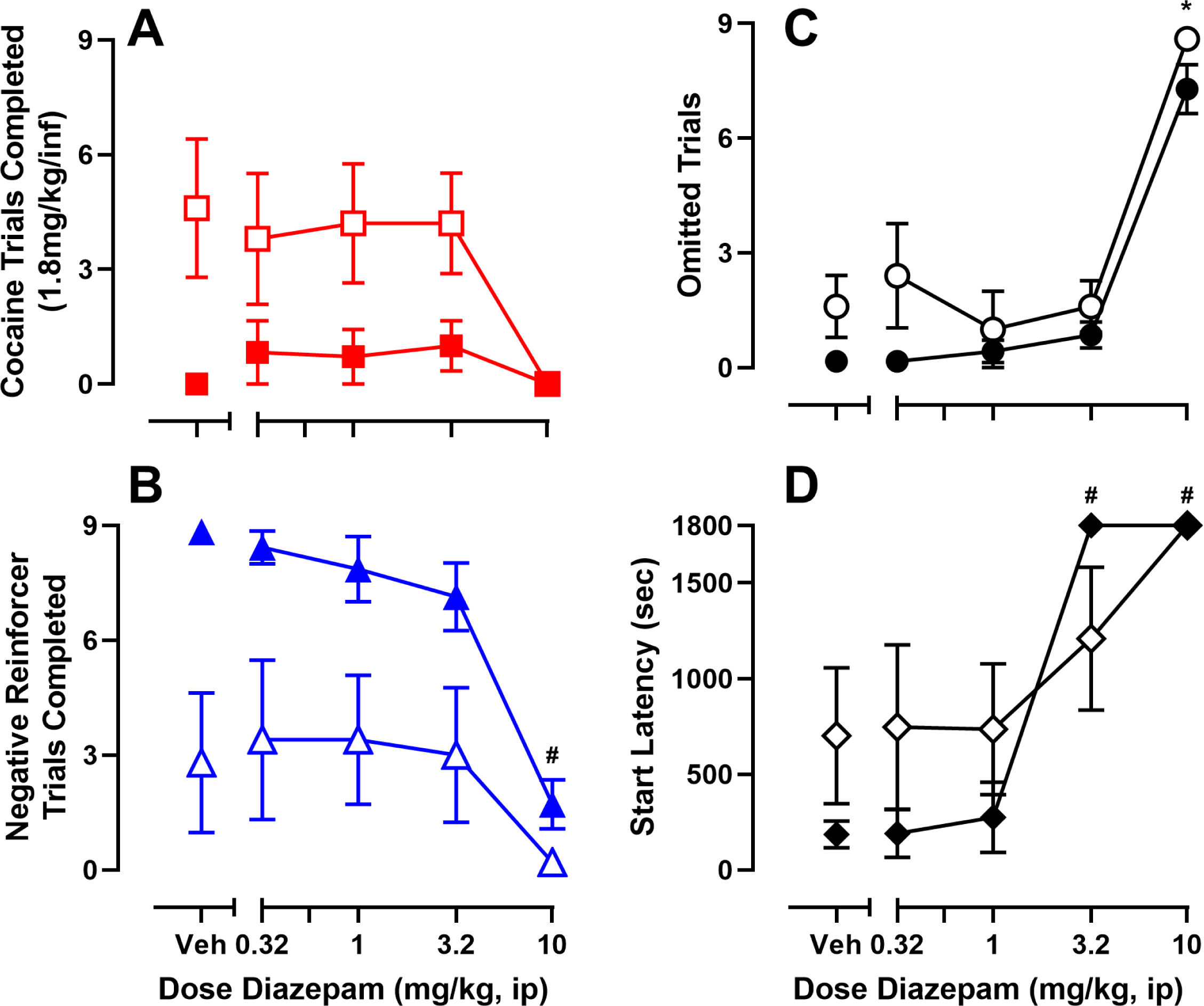
Effect of diazepam pretreatment (0.32 – 10 mg/kg, ip) on cocaine-vs-negative reinforcer choice. Abscissae: diazepam dose. (A) cocaine trials completed (B) negative reinforcer trials completed, (C) omitted trials, and (D) latency to respond for the first cocaine infusion. Filled and open symbols denote rats tested at the 0.7 mA (n = 7, 4F/3M) and 0.3 mA (n = 5, 2F/3M) shock condition, respectively. All points represent the mean ± SEM. Symbols represent significant (p < 0.05) comparisons: ^#^different from vehicle only in the 0.7mA shock group; * different from vehicle in both shock intensity groups.

## Discussion

The present study established a discrete-trial cocaine-vs-negative reinforcer choice procedure in male and female rats to test hypotheses related to drug-taking despite negative consequences in a choice context. Using this procedure, we determined the sensitivity of cocaine-vs-negative reinforcer choice to environmental and pharmacological manipulations that have been shown to impact cocaine-vs-positive reinforcer (e.g., food or social interaction) choice. There were three main findings. First, the presence of a concurrently available negative reinforcer robustly decreased cocaine self-administration up to 1.8 mg/kg/infusion. These results are consistent with prior studies showing that some concurrently available non-drug positive reinforcers, such as social interaction [26], or a saccharin solution [30] can robustly decrease cocaine-maintained behavior. Second, extended cocaine self-administration sessions resulted in high levels of cocaine intake but failed to promote behavioral reallocation towards cocaine and away from the negative reinforcer. These results do not support the hypothesis that extended cocaine self-administration leads to an addiction-like phenotype of increased drug-taking behavior despite adverse consequences in a choice context [9]. Finally, in contrast to the published single operant literature [31–34], acute diazepam pretreatment failed to alter cocaine-vs-negative reinforcer choice, further demonstrating that results from single operant drug self-administration studies have poor translation to preclinical drug-choice studies. Overall, the present results demonstrate that a concurrently available negative reinforcer, similar to a positive reinforcer, can attenuate addictive drug self-administration and highlights the importance of environmental context in preclinical models of drug addiction.

### Concurrently available negative reinforcement suppresses cocaine self-administration

Contrary to our hypothesis, a concurrently available negative reinforcer (i.e., escape/avoid shock) suppressed nearly all cocaine choices and there was minimal behavioral allocation between the two reinforcers across a range of cocaine doses and shock amplitudes. Previous cocaine-choice studies in both nonhuman primates and rats have demonstrated that behavioral allocation between cocaine and another positive reinforcer including both drug and nondrug reinforcers was sensitive to parametric manipulations of reinforcer magnitude and response requirement [24–28]. Furthermore, behavioral allocation in these cocaine-choice studies was sensitive to positive punishment with either electric shock [17] or IV histamine [16,19] such that behavior was reallocated away from the reinforcer paired with the punisher and towards the alternative, unpunished reinforcer. There is currently debate in the literature regarding whether the distinction between positive and negative reinforcement is empirically founded and involve distinctly different processes [35–38]. The present choice results support this distinction between positive and negative reinforcement and provide empirical evidence to support future research on both the quantitative and qualitative differences between positive and negative reinforcers.

One potential explanation for how positive and negative reinforcers are distinct could be within the framework of behavioral economics and the degree to which commodities including drugs as reinforcers function as substitutes, complements, or independents (for review, see [39]). For example, numerous studies have shown that increasing the magnitude of the food reinforcer promotes behavioral allocation towards food choice and away from cocaine choice. Inversely, increasing the available dose of cocaine promotes a reallocation of behavior towards cocaine choice [26,27,40]. This same relationship also holds for response requirement manipulations [25,27]. In behavioral economic terms, food and cocaine when concurrently available as positive reinforcers could be considered economic substitutes [39]. In the present study, behavioral allocation between cocaine and a negative reinforcer was minimally sensitive to reinforcer magnitude and response requirement manipulations, suggesting that cocaine and the negative reinforcer were not economic substitutes. Rather, cocaine and the negative reinforcer in this choice context appeared to be economic independents [39]. Even at small shock amplitudes, rats omitted trials despite receiving electric shock rather than complete the response requirement on the cocaine-associated lever. Overall, the present results suggest that although negative reinforcement may decrease cocaine self-administration, it is not an economic substitute for cocaine. Considering this, treatment strategies that utilize a combination of both nondrug positive and negative reinforcement contingencies might be most effective for treatment of cocaine use disorder.

### Extended cocaine access failed to increase cocaine-vs-negative reinforcer choice

Increased drug availability through extended access drug self-administration conditions is one common preclinical method to achieve high levels of cocaine intake [41]. Cocaine self-administration under extended access conditions is hypothesized to be associated with the transition to “compulsive” drug use [41,42], often operationalized as continued drug-taking behavior despite adverse consequences [9,10,45–47]. Preclinical studies have demonstrated that extended cocaine access can lead to a decreased sensitivity to shock-associated punishment of cocaine reinforcement in a subset of rats [9,10,43]. The present results show that male and female rats achieved high levels of cocaine intake during extended access sessions; however, in contrast to previous findings and despite the use of similar shock magnitudes to those used in previous studies [9], extended cocaine access failed to increase cocaine-vs-negative reinforcer choice. Furthermore, out of 20 total rats, only a single rat completed more cocaine trials during the choice session after the extended cocaine access sessions, indicating an overall absence of this “compulsive” phenotype. The present results are consistent with and extend previous cocaine-vs-food choice studies in rhesus monkeys demonstrating extended access cocaine self-administration failed to increase cocaine choice [44]. Overall, the present results suggest that the behavior of cocaine-taking despite adverse consequences may be driven more by a lack of alternative reinforcers, rather than an “compulsive phenotype” [45].

### Acute diazepam did not increase cocaine-vs-negative reinforcer choice

Under single operant behavioral conditions, benzodiazepine administration increases punished responding maintained by multiple reinforcer types [31–34] including addictive drugs [46]. Therefore, we hypothesized that acute diazepam pretreatment would increase cocaine choice in the cocaine-vs-negative reinforcer choice procedure. However, diazepam pretreatment failed to increase cocaine-vs-negative reinforcer choice, suggesting that benzodiazepines may only increase punished responding under single operant contingencies. Diazepam administration before a cocaine-vs-food choice session in rats has been shown to decrease cocaine choice and increase food choice in the absence of any punishment contingencies [47]. Overall, these results add to the current literature highlighting the importance of incorporating alternative reinforcers into drug self-administration models and that results from single operant drug self-administration procedures do not always translate to drug self-administration procedures that include concurrent availability of another reinforcer. Future experiments should evaluate diazepam pretreatment effects on punished drug-vs-alternative positive reinforcer choice procedures.

## Acknowledgments

We appreciate the technical assistance of Michelle Arriaga during the training procedures.

## Author Contributions

**Madison Marcus:** Writing – Original Draft, Writing – Review and Editing, Methodology, Software, Formal analysis, Investigation, Data Curation, Data Visualization. **Matthew Banks:** Formal analysis, Writing – Review and Editing, Visualization, Conceptualization, Supervision, Funding Acquisition.

