## Supplemental Materials for "A concurrently available negative reinforcer robustly decreases cocaine self-administration in male and female rats"

| | Negative Reinforcement (Escape) | | | Cocaine Self-Administration | | |
|---|---|---|---|---|---|---|
| | *Rats in group* | *Rats to meet criteria* | *% to meet criteria* | *Rats in group* | *Rats to meet criteria* | *% to meet criteria* |
| NR First | 12<br>(6M /6F) | 11<br>(6 M /5 F) | 92 | 10<br>(5 M /5 F) | 10<br>(5 M/ 5 F) | 100 |
| Cocaine First | 10<br>(5 M /5 F) | 10<br>(5 M /5 F) | 100 | 10<br>(5 M /5 F) | 9<br>(4 M/ 5 F) | 90 |
| Food First | 4<br>(2M/2F) | 4<br>(2M/2F) | 100 | 3<br>(2 M /1 F) | 3<br>(2 M /1 F) | 100 |

Table S1. Acquisition success rates for negative reinforcement and cocaine self-administration training according to first training experience.

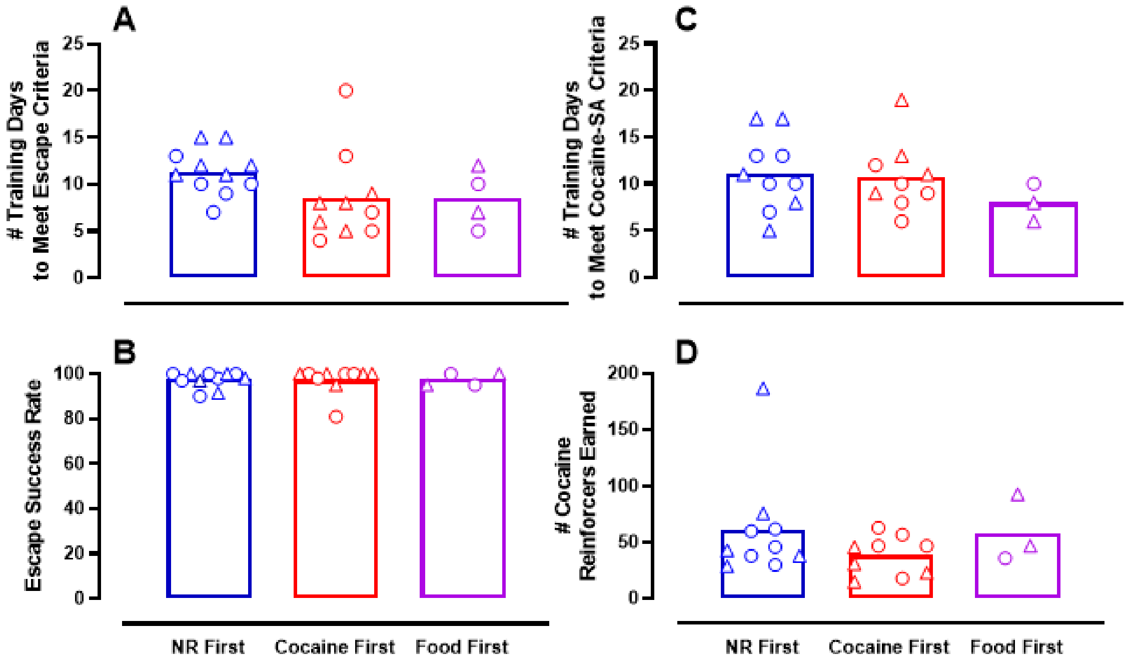


Figure S1. No effect of first training experience on acquisition of escape (A-B) or cocaine self-administration responding (C-D). Abscissae: first training experience. Points represent individual subject data from the final day of training. Only rats who met acquisition criteria are included. Triangles and circles represent males and females, respectively. Bars represent group mean.

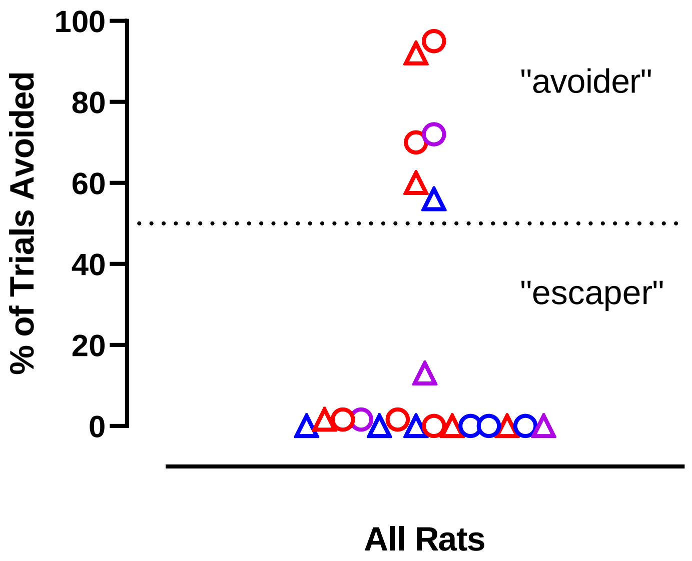


Figure S2. Individual avoidance behavior during negative reinforcement training. Avoidance of ≥ 50% of trials (dashed line) resulted in “Avoider” classification, ≤ 50% avoidance resulted in “Escaper” classification. Points represent individual subject data from the final day of training. Colors correspond to first training experience of the subject. Blue: negative reinforcement first; Red: cocaine first; Purple: food first. Triangles and circles represent males and females, respectively.

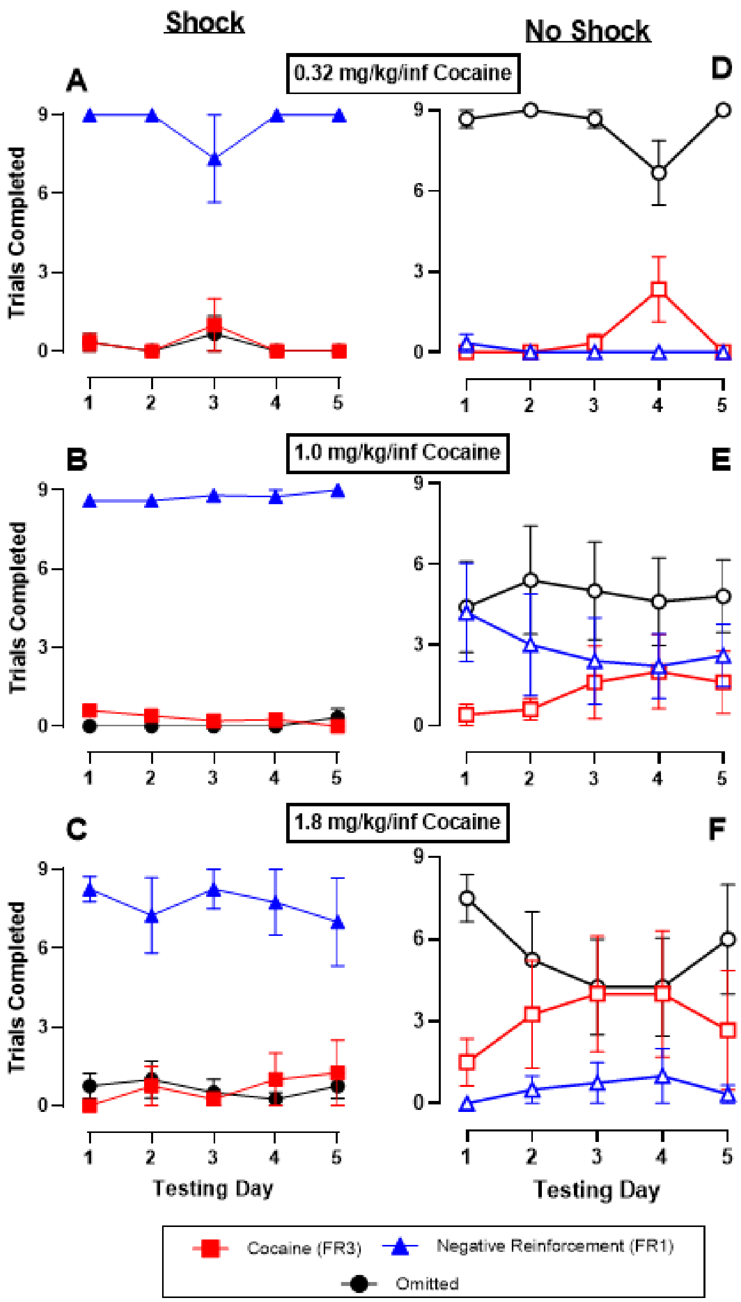


Figure S3. Cocaine-vs-negative reinforcer choice across five consecutive testing days for three cocaine doses. Abscissae: testing day. Ordinates: number of trials completed for cocaine, negative reinforcement, or omitted. Shaded and open symbols represent shock and no shock conditions, respectively. Points represent mean ± SEM. (A, D) n = 3 (2F/1M); (B, E) n = 5 (4F / 1M); (C, F) n = 4 (2F / 2M)

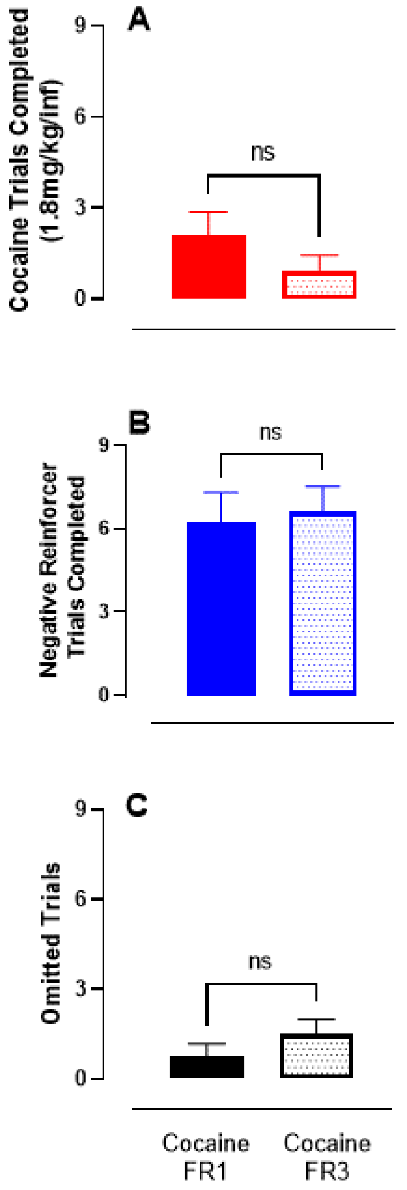


Figure S4. No effect of cocaine response requirement on cocaine-vs-negative reinforcer choice. Trials completed for (A) cocaine, (B) negative reinforcement, and (C) omitted when cocaine is on an FR1 (solid bars) and FR3 (patterned bars) schedule of reinforcement. Rats chose between cocaine (1.8 mg/kg/inf) and a negative reinforcer (escape or avoidance of 0.7 mA shock, FR1) for five days at each response requirement and the results of Day 5 are shown. Bars represent the group mean ± SEM. n = 10 (4F/6M). ns = non-significant T-test

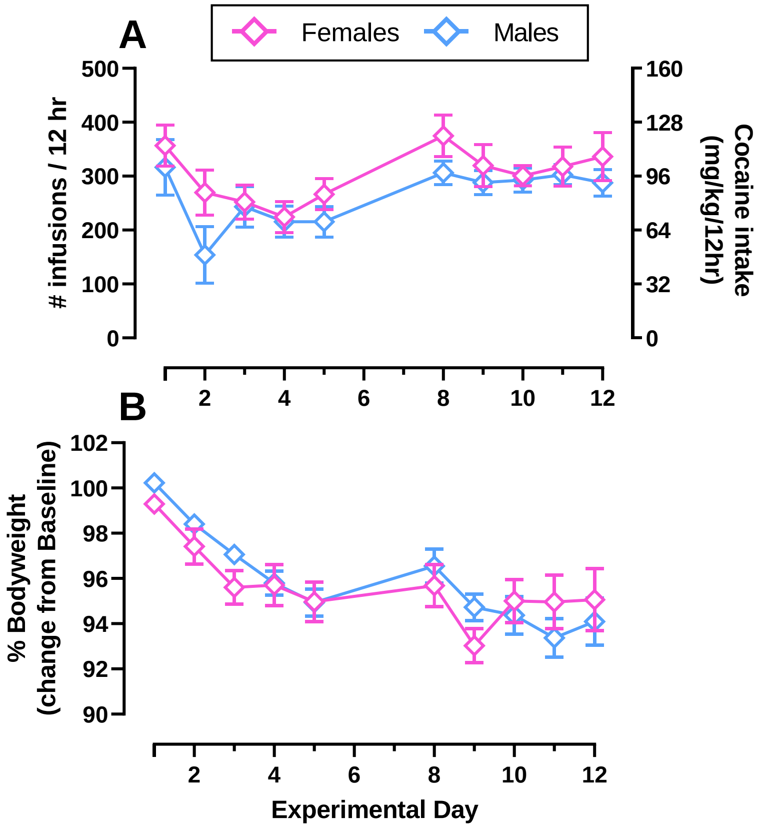


Figure S5. No sex differences detected under extended access conditions. (A) number of cocaine infusions self-administered over the 12-hr period (0.32 mg/kg/inf, FR3 / 10-s time-out) (B) bodyweight change from baseline. Points represent the mean ± SEM. n = 20 (10F/10M).

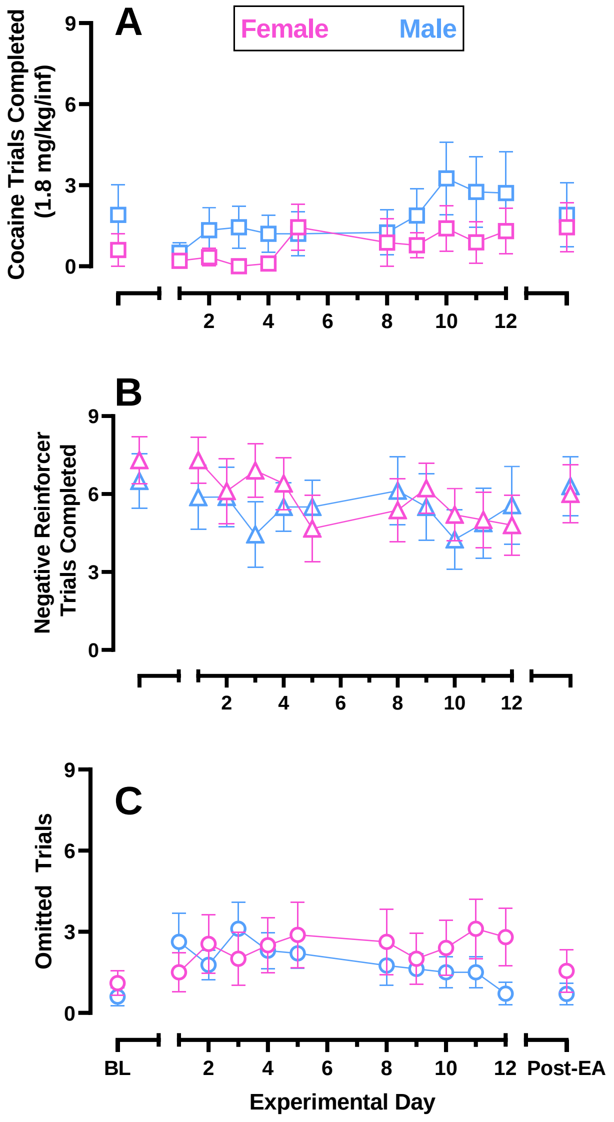


Figure S6. No sex differences detected in cocaine-vs-negative reinforcer choice under extended access cocaine conditions. Abscissae: experimental day. Ordinates: number of trials completed for (A) cocaine (1.8 mg/kg/inf, FR3), (B) negative reinforcement (0.7 mA, FR1), or (C) omitted. Pink and blue represent female and male subjects, respectively. Shapes correspond to trial type. Baseline (BL) is the Friday prior to initiating extended cocaine access. Post-EA is seven days after terminating extended cocaine access. Points represent the mean ± SEM. n = 20 (10F/10M).

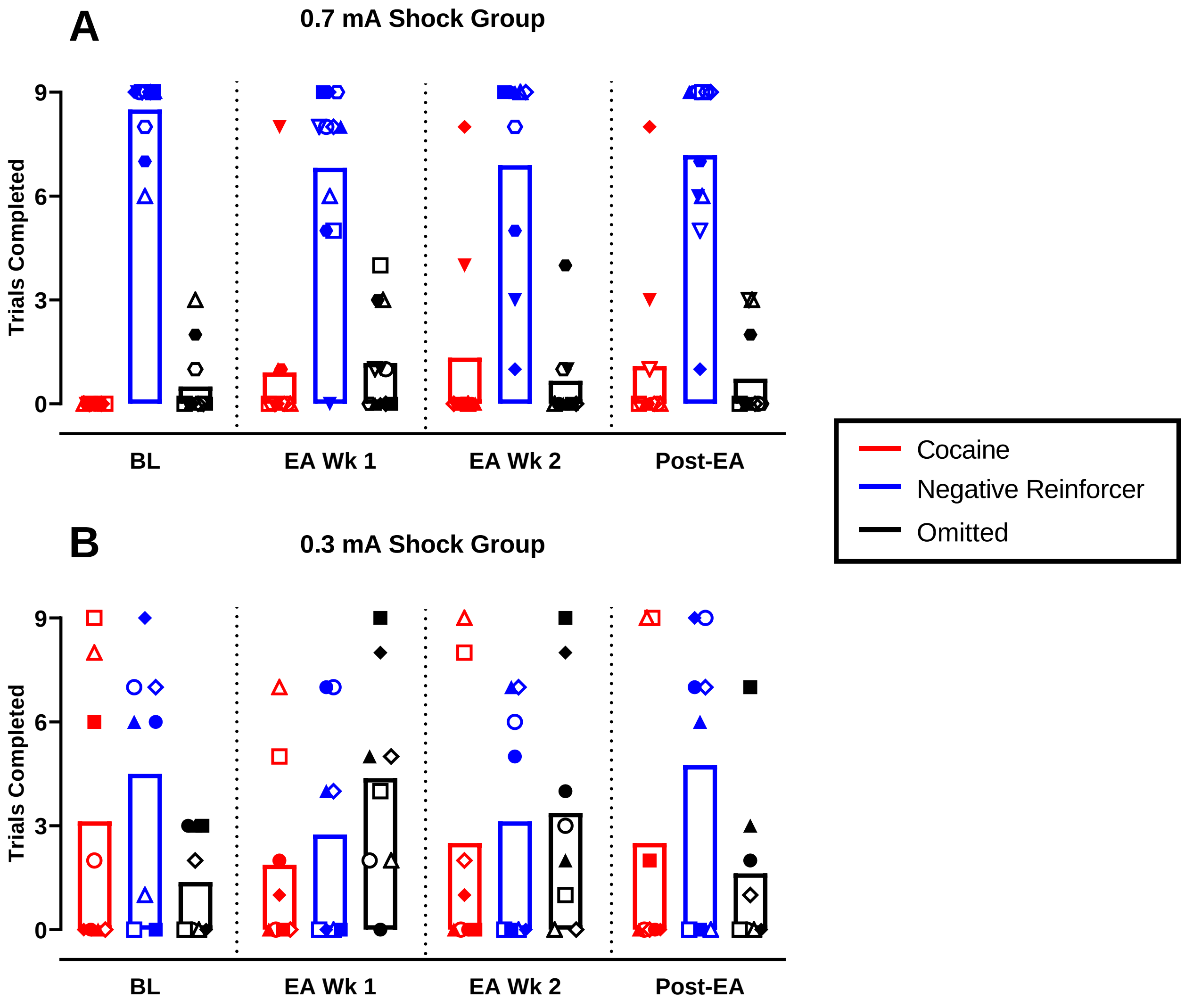


Figure S7. Cocaine-vs-negative reinforcer choice of individual rats during extended cocaine access conditions. In the 0.7 mA condition, only one rat (filled diamond) started taking cocaine after extended cocaine access. In the 0.3 mA condition, zero rats started taking cocaine after extended cocaine access. Baseline (BL) is the Friday prior to initiating extended cocaine access, EA Wk1 and 2 are the Friday of the first and second week of extended cocaine access, respectively. Post-EA is seven days after terminating extended cocaine access. Ordinates: trials completed for cocaine (red), negative reinforcement (blue), or omitted (black). Bars represent the group mean; each symbol corresponds to an individual subject. Filled and open symbols represent female and male subjects, respectively. 0.7 mA shock group n = 12 (6F/6M); 0.3 mA shock group n = 8 (4F/4M).
